# Structural and biochemical characterization on the cognate and heterologous interactions of the MazEF-mt9 TA system

**DOI:** 10.1101/432989

**Authors:** Ran Chen, Jie Tu, Yaoju Tan, Xingshan Cai, Chengwen Yang, Xiangyu Deng, Biyi Su, Shangming Ma, Xin Liu, Pinyun Ma, Chaochao Du, Wei Xie

## Abstract

The toxin-antitoxin (TA) modules widely exist in bacteria, and their activities are associated with the persister phenotype of the pathogen *Mycobacterium tuberculosis* (*M. tb*). *M. tb* causes Tuberculosis, a contagious and severe airborne disease. There are ten MazEF TA systems in *M. tb*, which play important roles in stress adaptation. How the antitoxins antagonize toxins in *M. tb* or how the ten TA systems crosstalk to each other are of interests, but the detailed molecular mechanisms are largely unclear. MazEF-mt9 is a unique member among the MazEF families due to its tRNase activity, which is usually carried out by the VapC family toxins. Here we present the cocrystal structure of the MazEF-mt9 complex at 2.7 Å. By characterizing the association mode between the TA pairs through various characterization techniques, we found that MazF-mt9 not only bound its cognate antitoxin, but also the non-cognate antitoxin MazE-mt1, a phenomenon that could be also observed *in vivo*. Based on our structural and biochemical work, we proposed that the cognate and heterologous interactions among different TA systems work together to relieve MazF-mt9’s toxicity to *M. tb* cells, which may facilitate their adaptation to the stressful conditions encountered during host infection.

**IMPORTANCE:** Tuberculosis (TB) is one of the most severe contagious diseases. Caused by *Mycobacterium tuberculosis* (*M. tb*), it poses a serious threat to human health. Additionally, TB is difficult to cure because of the multipledrug-resistant (MDR) and extensively drug-resistant (XDR) *M. tb* strains. Toxin-antitoxin (TA) systems have been discovered to widely exist in prokaryotic organisms with diverse roles, normally composed of a pair of molecules that antagonize each other. *M. tb* has ten MazEF systems, and some of them have been proved to be directly associated with the genesis of persisters and drug-resistance of *M. tb*. We here report the MazEF-mt9 complex structure, and thoroughly characterized the interactions between MazF-mt9 with MazEs within or outside the MazEF-mt9 family. Our study not only revealed the crosstalks between TA families and its significance to *M. tb* survival but also offers insights into potential anti-TB drug design.

---

The toxin-antitoxin (TA) modules widely exist in bacteria, whose activities have been found to be associated with the persister phenotype (1, 2). In most cases, the toxins act upon specific cellular processes, while the corresponding antitoxins antagonize their activities (3, 4).

In a typical TA system, the two genes are located adjacently to each other and share the same operon. As a result, their transcription and translation are usually coupled to ensure proper stoichiometric quantities of both molecules (5). The toxins are invariantly proteins and their molecular weights are usually smaller than 10 kDa (4). The cores of these proteins typically feature a β-sheet (6–8). On the other hand, the antitoxins could either be proteins or RNAs, which neutralize the functions of toxins. Due to their distinct functions, toxins and antitoxins have different life spans. Generally speaking, toxins are much more stable than their antitoxins and therefore can exist for longer periods (9–11). Antitoxins as proteins usually have loose and flexible structures, and they are prone to degradation. When cells undergo environmental stresses, the balance between toxins and antitoxins is broken. Antitoxins will be degraded and toxins thus released act on their substrates, leading the cells to the bacteriostasis stage (5, 12). Depending on how antitoxins regulate the activities of toxins, TA systems are divided into six different classes (13–15).

The *Mycobacterium tuberculosis* (*M. tb*) genome contains more than 90 putative toxin-antitoxin operons, including ten MazEF systems (designated as mt1–10 respectively) (16, 17). It has been found that overexpression of the *maz*F-mt1, *maz*F-mt3, *maz*F-mt6 and *maz*F-mt9 genes in the *Mycobacterium bovis* BCG or *M. tb* H37Rv strains induced bacteriostasis (18, 19). In addition, their respective encoded products, *maz*F-mt1, *maz*F-mt3 and *maz*F-mt6 also responded to a variety of stresses (18). MazFs are generally RNA endonucleases, acting on mRNA or rRNA. Using an RNA-seq approach, Schifano *et al.* identified that the principal substrates for MazF-mt9 were two tRNAs-tRNA^Pro(GGG)^ and tRNA^Lys(UUU)^, and the cleavage occurred within a UUU motif in either the D-or the anticodon loop (19).

The structure of *Escherichia coli* MazF (EcMazF) in complex with MazE was determined in 2003 (PDB 1UB4) (20). A linear MazF2-MazE2-MazF2 heterohexamer was formed, where the MazE homodimer occupies the middle and interacts with the two flanking MazF homodimers. Simanshu *et al.* solved the structure of the MazEF complex from *Bacillus subtilis* (PDB 4ME7) (21). The BsMazE structure is dissimilar to that of the EcMazEF complex (PDB 1UB4) (20), suggesting possible shuffling of the antitoxin folds among different TA systems. They also reported the structure of BsMazF complexed with a 9-nt RNA fragment, which binds to the toxin’s interfacial groove, explaining how the antitoxin inhibits the mRNase activity of BsMazF (PDB 4MDX) (21). Recently Hoffer *et al.* reported the crystal structure of *M. tb* MazF-mt6 (PDB 5UCT), which cleaves 23S rRNA Helix 70 to inhibit protein synthesis (22). MazF-mt6 adopts a PemK-like fold but lacks an elongated β1-β2 linker typically seen in other MazF members. The structures of MazF-mt7 and the MazEF-mt7 complex were determined immediately after the work on MazF-mt6, and the respective interaction sites of the MazF-mt7 toxin with an RNA substrate and a new EDF homolog were mapped using NMR spectroscopy (PDBs 5XE2 and 5XE3) (23). In addition, the structures of MazF-mt1, MazF-mt3 and their variants in complex with RNA or DNA were also released recently (PDBs 5HJZ, 5HK0, 5HK3 and 5HKC), but the related work was not published.

Previously we determined the crystal structure of *M. tb* MazF-mt9 in its apo form and investigated its biochemical properties (24). Structural analysis revealed a highly positively charged dimer interface formed by MazF and we proposed that this deeper and wider interface should be the structural basis for MazF to recognize and cleave tRNA substrates. To further elucidate the molecular basis on which the antitoxin MazE inhibits MazF’s function, we solved the crystal structure of the *M. tb* MazEF-mt9 complex at 2.7 Å and characterized the association mode between the toxin-antitoxin pair. We found that MazF-mt9 not only bound to its cognate antitoxin, it also bound non-cognate antitoxin MazE-mt1. Our studies suggested that toxins and antitoxins from different families could cross talk, which may serve as a mechanism to offset the toxicity to *M. tb* cells.

## RESULTS

### Overall structure of the MazEF-mt9 complex

To better understand how the two proteins of the MazEF-mt9 family interact with each other, we solved the structure of the MazEF-mt9 complex. Our initial attempts to obtain the binary complex through co-expression were unsuccessful because the two proteins failed to form a stable complex, possibly due to the interference from the 8 × His-tag present at the N-terminus of MazE-mt9. Therefore, the two proteins were individually expressed and incubated together to allow their association and subsequent crystallization after the removal of the tag. In the resulting cocrystal structure, two MazEs and MazFs were present in the asymmetric unit. They each formed their own homodimers first, and then a hetero-tetramer (Fig. 1A). While the two MazF monomers were very similar in structure (a 0.21 Å RMSD over 112 Cαs, Fig. 1B), α3’ in one of the MazE-mt9 monomers was degraded (chain D) but its counterpart in the other MazE-mt9 monomer was intact. The two MazF-mt9 monomers were almost completely resolved except for a small fragment close to the N-terminus (A17-G23 for chain A and A17-P22 for chain B respectively, Table 1). The more structured MazE-mt9 monomer (chain C) consists of a β-strand and three α-helices, forming the “head” (β1) and “tail” (α1-α3) domains respectively. The starting β-strand forms an antiparallel sheet with its equivalent strand from the other monomer. α2 is the longest helix in MazE-mt9 and these two helices from the two subunits cross over at an angle of ~120°. Chain C of the MazE-mt9 dimer binds across the dimer interface formed by MazFs, and then it sharply bends at its α2-α3 junction, forming an angle ~90°. α3’ was not well defined in our density map and it was probably degraded even before crystallization. Additionally, MazFs form a dimer that is almost identical in shape to that of its apo form (PDB 5WYG), with a 0.4 Å over 112 Cαs, indicating that the binding of MazE-mt9 caused minimal structural rearrangements of the nuclease (Fig. 1C).

**FIG 1.**
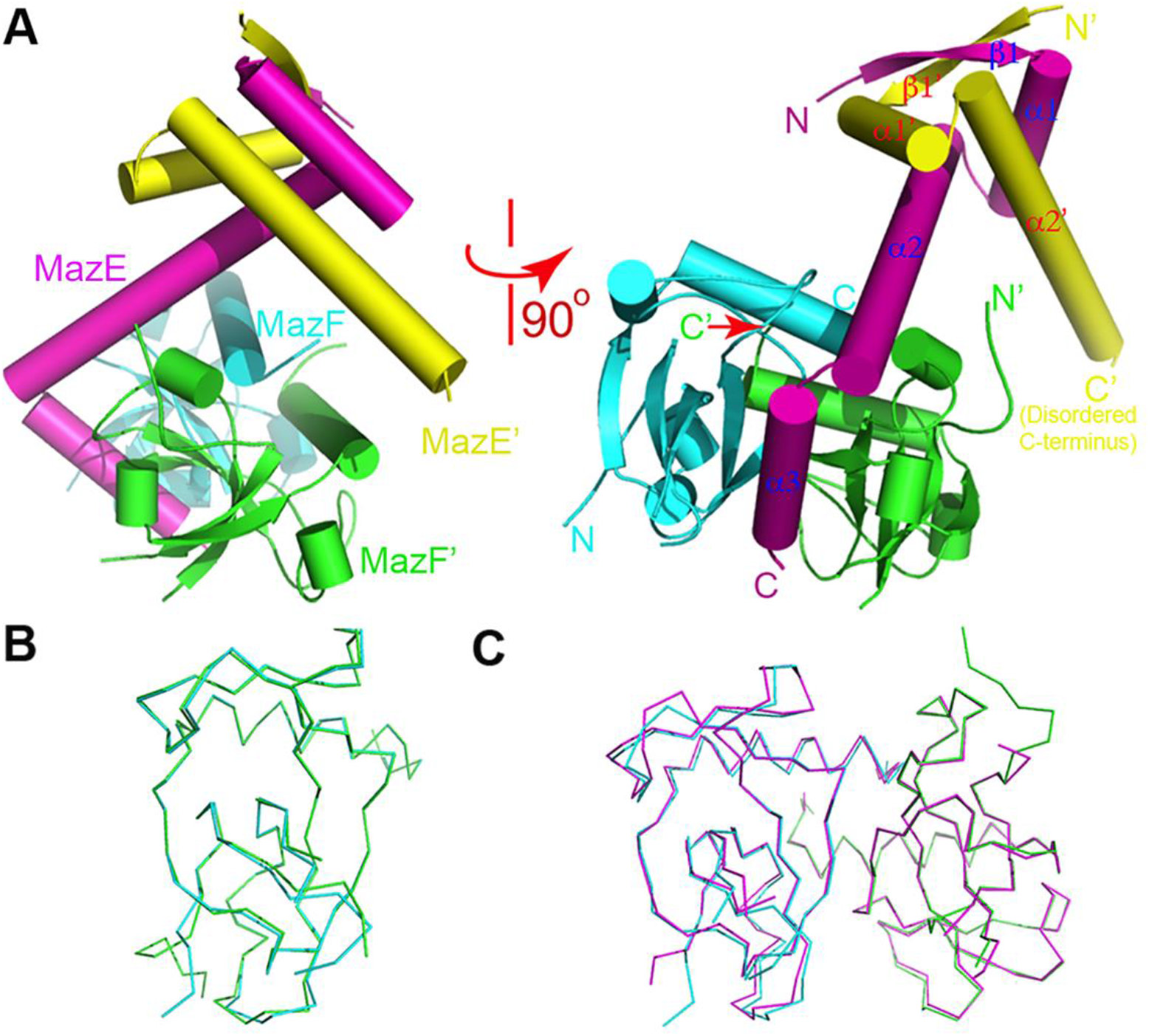
The overall view of the MazEF-mt9 complex and structure comparison. (A) The structure shown in the cylinder rendition in two orthogonal views. The two MazF-mt9 monomers are colored cyan and green respectively while the two MazE monomers are in magenta and yellow respectively. The N-and C-termini are indicated and the secondary structural elements of MazE-mt9 are labeled. (B) The structural alignment of the Cα traces of the two MazF-mt9 monomers in the complex. (C) The structural alignment of the Cα traces of MazF-mt9 before (in green/cyan) and after (in magenta) the binding of MazE-mt9 (MazE-mt9 was not shown for clarity in both (B) and (C).

**TABLE 1.**
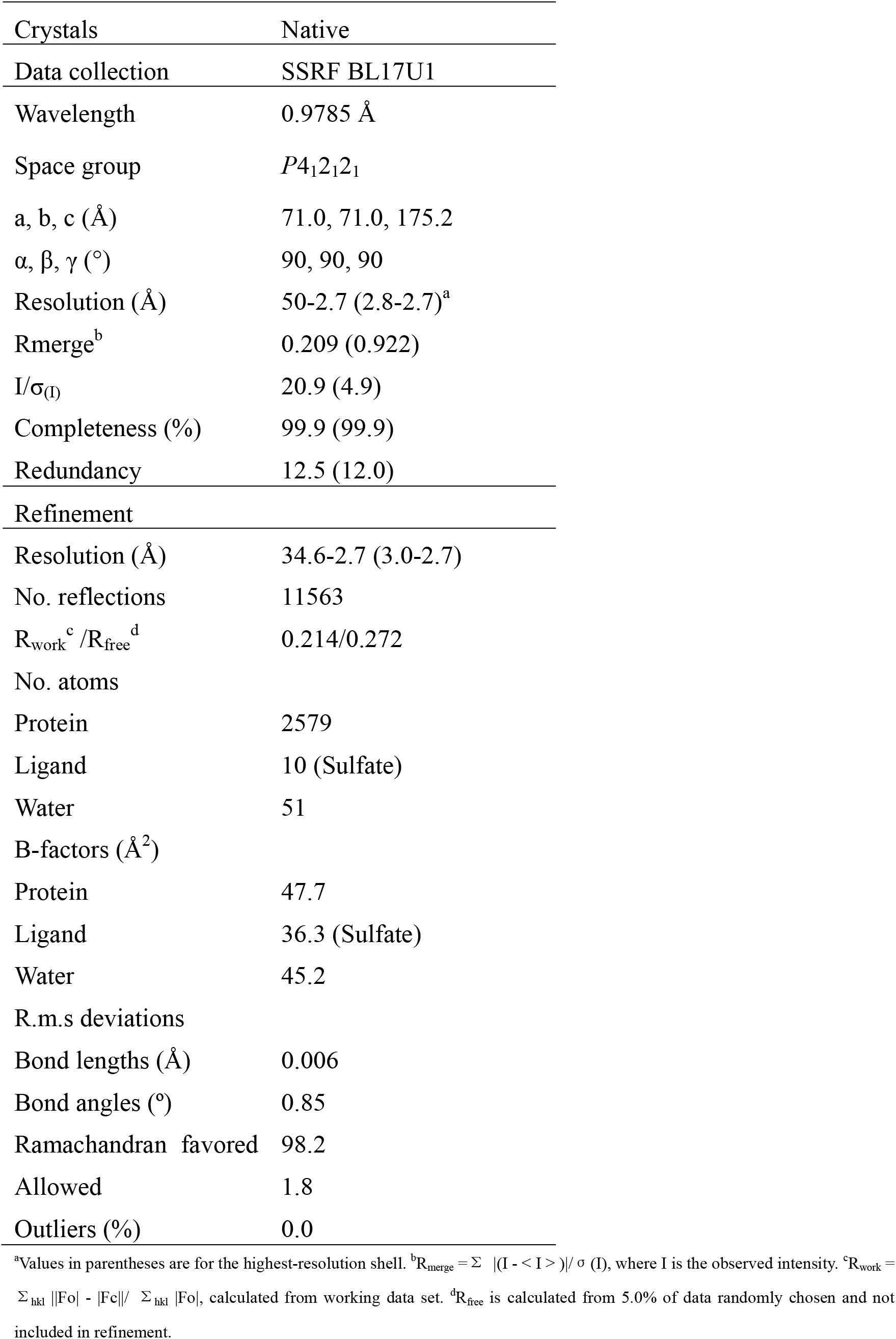
Data collection and refinement statistics.

The binding mode of the *M. tb* MazEF-mt9 complex is closely related to that of *B. subtilis* MazEF (BsMazEF, PDB 4ME7). Superimposition of the two complexes revealed that the MazF dimer structures between the two species are very similar (not shown), but the MazE dimers are somewhat different (Fig. 2A). Particularly, the β1-α1 loop is longer in *B. subtilis* than its counterpart in *M. tb* (4 vs. 2 residues), causing the residues in the α2-helix generally to translate ~5–8 Å. In addition, the angle formed by the α2-and α3-helices in BsMazE is ~120 ° while it is nearly a right angle in *M. tb*, and the α3-helix is kinked in the former structure to make more contacts with BsMazF. Interestingly, residues A17-G23 are invisible in *M. tb* MazF, which are also disordered in BsMazF. Despite the dissimilarities in MazE, the general binding mode between the toxin-antitoxin is conserved, in which the antitoxin molecules occupy the interface of the toxin dimers. Our structure also resembles that of the *M. tb* MazEF-mt7 complex (PDB 5XE3), but the starting fragment in MazE-mt7 does not form an antiparallel sheet with the other monomer, and the α2 helix is more straight than its counterpart in MazE-mt9 (Fig. 2B).

**FIG 2.**
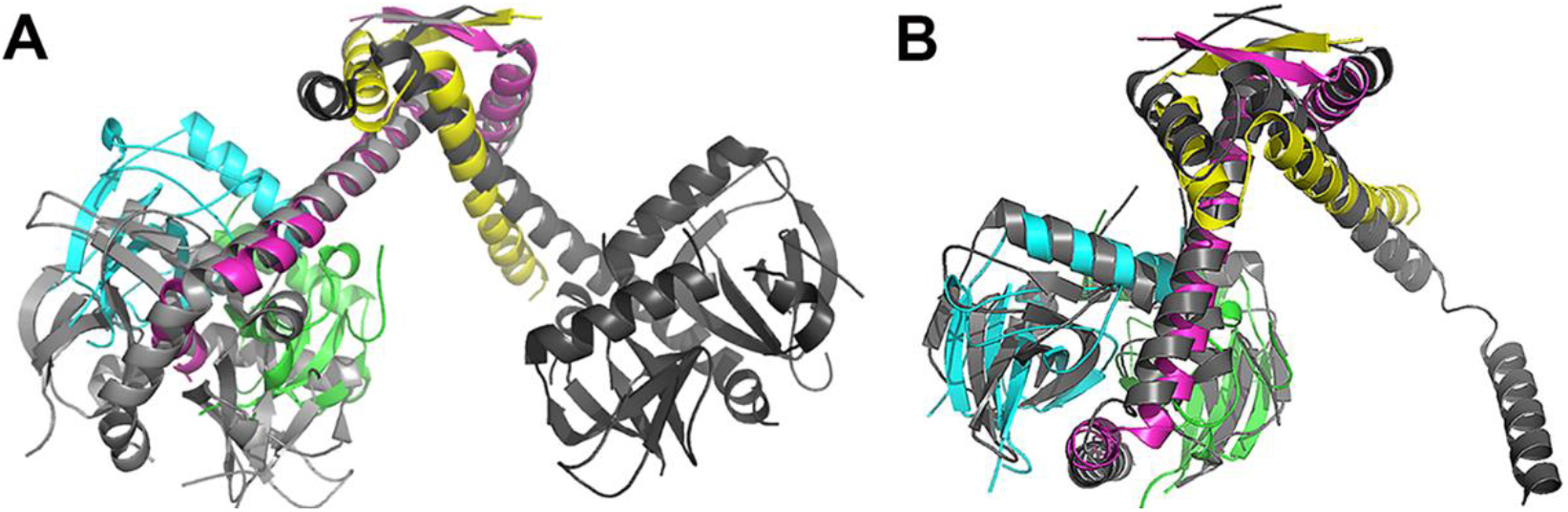
Structure comparison of the MazEF-mt9 complex with closely related complexes. (A) The structural superimposition of the *M. tb* MazEF-mt9 and BsMazEF-mt9 (PDB 4ME7) complexes, with the latter showing in gray. (B) The structural superimposition of the *M. tb* MazEF-mt9 and MazEF-mt7 (PDB 5XE3) complexes, with the latter showing in gray. Chains C and D of BsMazF and MazEF-mt7 were omitted for clarity.

### Characterization of the affinity between the MazE/F-mt9 complex

There are extensive protein-protein interactions between MazE-mt9 and MazF-mt9, including hydrogen bonds, salt bridges and hydrophobic interactions, to fix the MazE monomer into the binding groove of MazF-mt9. The interactions were listed in Table S1, and the total buried surface was calculated to be 1209 Å^2^. These interactions are mainly distributed in two areas in MazE-mt9: the last helix α3 and the C-terminal half of α2. Specifically, the negatively charged dipeptide D63E64, makes a salt bridge and a hydrogen bond with the positively charged residues R81 and R26 of the MazF-mt9 dimers (Fig. 3A). D74 makes more complex interactions with the toxin, adding five extra hydrogen bonds that involve some backbone contacts. On the other hand, the interactions concerning α2 mainly concentrate in two sites: the I44-E50 and the E55 sites. The former forms a network of mixed interactions with MazF-mt9: while the dipeptide E48R49 forms hydrogen bonds with T57, R60 from one MazF-mt9 monomer, and with G-3’ (resulted from the protease digestion site, and primes indicates another monomer) and E35’ from another, the I44F45 dipeptide forms hydrophobic interactions with L59, L109, V33’ and L36’ (Fig. 3B). In contrast, E55 mainly makes salt bridges and hydrogen bonding interactions (Fig. 3C). To assess the contribution of each interaction to the stability of the complex, we selectively mutated key residues from MazE-mt9 and tested their impacts on the growth rates of their expression host *Escherichia coli*, after induction of the TA systems (the toxins included the wild type (WT) and single mutants). As a control experiment, we first checked the *E. coli* growth after the induction of *maz*F-mt9 alone. The expression of MazF-mt9 nearly brought a complete stop to the bacterial growth, with the OD_600_ curve plateauing at 0.6 (Fig. 4A). However, when MazF-mt9 was co-expressed with the WT or the MazE-mt9 mutants (including multiple mutants), *E. coli* showed similar growth rates to that of the strain transformed with an empty vector (which had a doubling time less than 30 min), but elevated rates than that of the WT-transformed strain in suspension cultures (Fig. 4A and B). These results indicated that MazE-mt9 bound so tightly to MazF-mt9 that mere single mutations would not disrupt their interactions. Consequently, the association of MazE-mt9 with MazF-mt9 protected the host tRNAs from MazF-mt9 degradation. The growth of *E. coli* colonies on LB-agar plates showed a similar trend (Fig. 4C and D). These results were also in agreement with previous observation by Simanshu *et al*. that single mutations of these residues alone were not sufficient to abolish the interactions between BsMazE and BsMazF (21).

**FIG 3.**
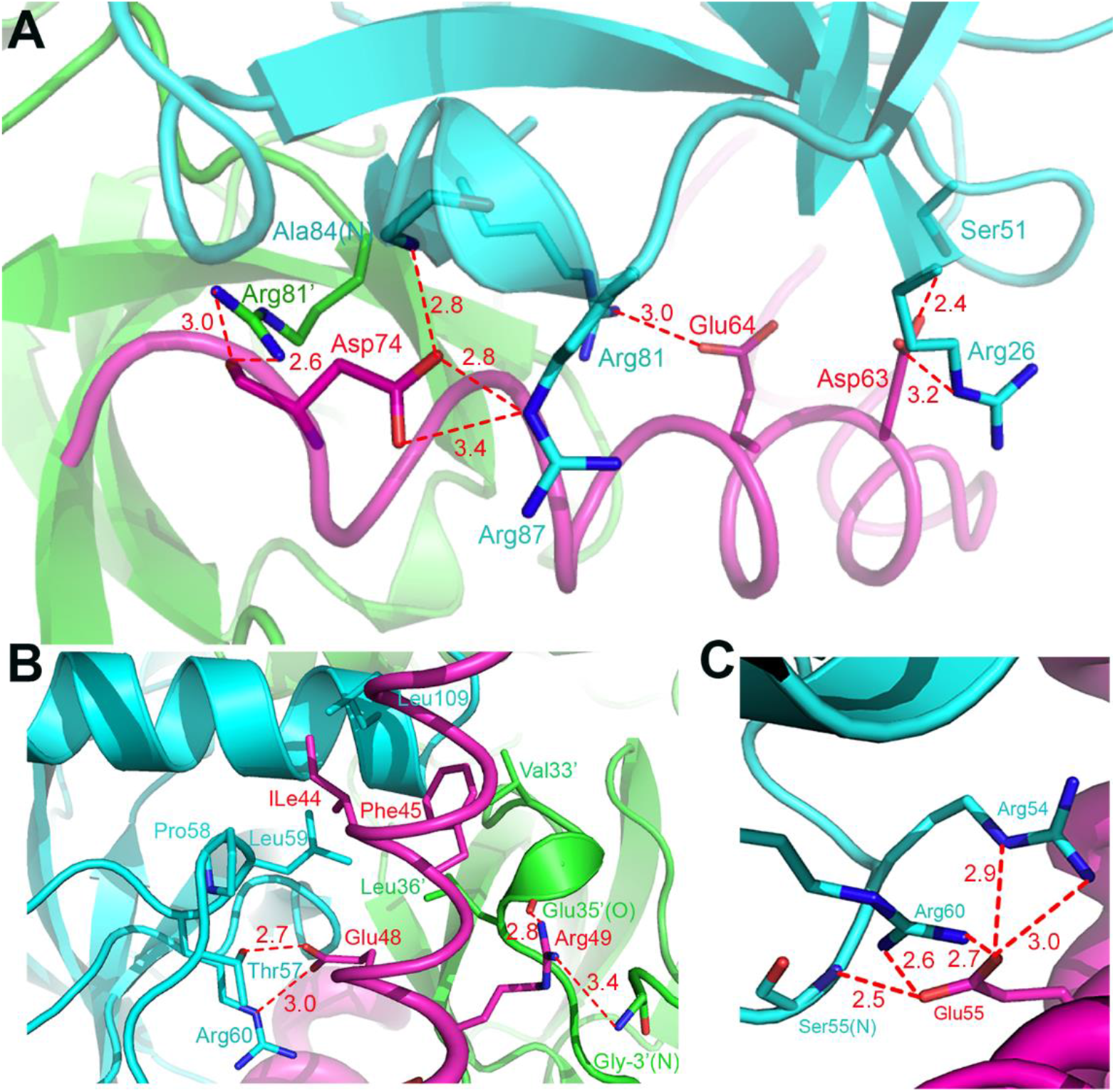
The detailed interactions between the MazEF-mt9 TA pairs. The hydrogen bonds are shown by the dashed lines and the distances are shown near the lines. (A) The interactions between MazE α3 (magenta) and the MazF dimer (cyan and green). Primes indicate residues from a different monomer. (B) The detailed interactions between MazE-mt9 around the I44-E50 fragment (magenta) and the MazF dimer (cyan and green). (C) The detailed interactions between residue E55 from MazE-mt9 (magenta) and the MazF dimer (cyan and green).

**FIG 4.**
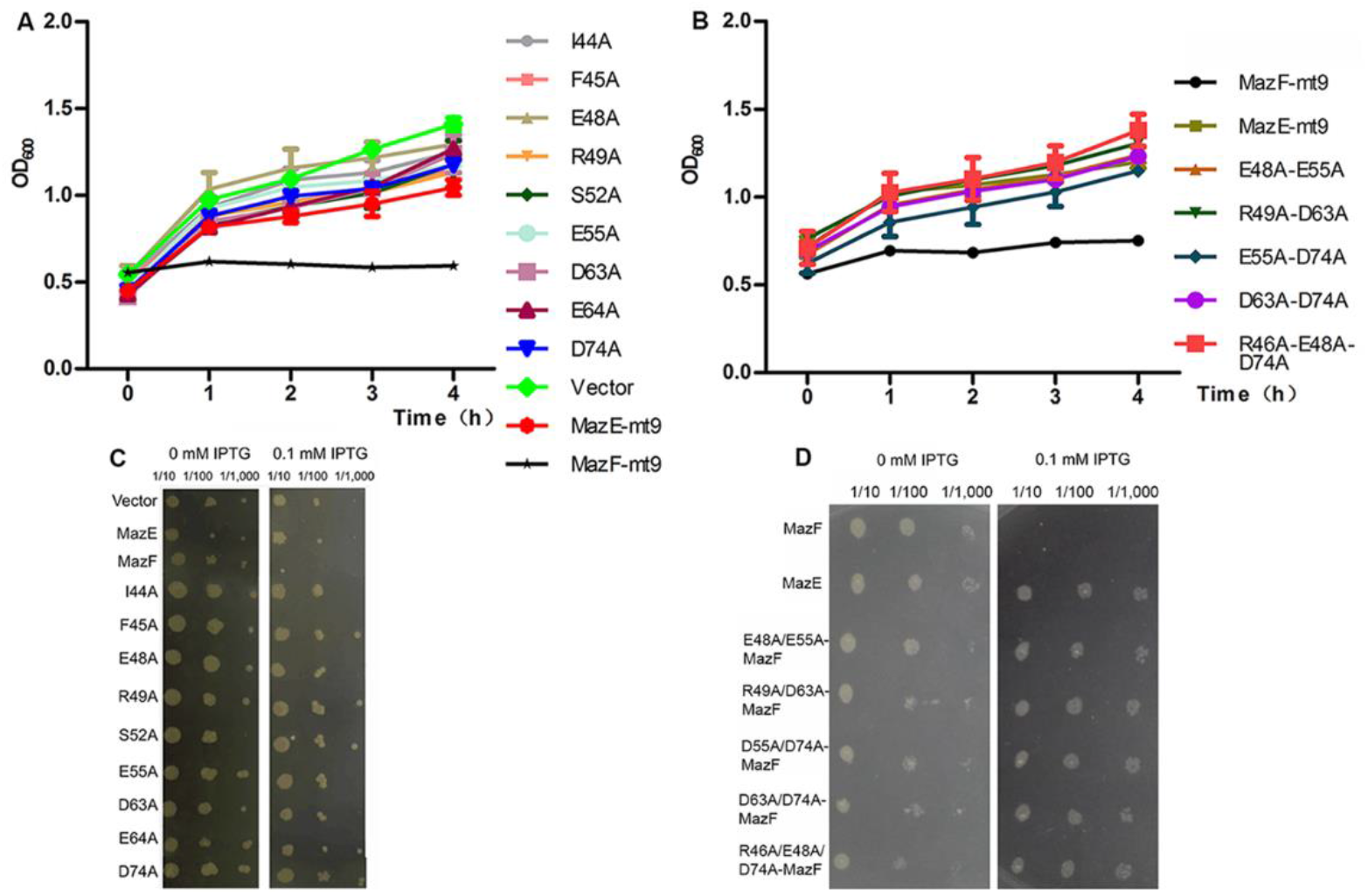
The influence of the MazEF-mt9 TA system on the *E. coli* host growth. (A-B) *E. coli* growth curves after the induction of the WT *maz*F or the co-expressed *maz*EF genes including single (A) or multiple (B) mutants of MazE, and the empty vector. (C-D) The growth of the *E. coli* colonies carrying the WT *maz*F or the co-expressed *maz*EF genes with different MazE mutations as tested in (A-B) on LB-agar plates.

To examine the subtle changes of the suboptimal association of the MazF-MazE pairs caused by mutations, individual MazE-mt9 mutants were expressed and their affinities to WT MazF-mt9 were evaluated indirectly by the cleavage activities of MazF-mt9 toward tRNA in the presence of the mutants. The addition of MazE-mt9 to MazF-mt9 at the 1:2 molar ratio resulted in 86% activity loss of cleavage activity (Fig. 5A and B). Similarly, pre-incubation of different single mutants of MazE-mt9 with MazF-mt9 also inhibited the activity of MazF-mt9, but to lesser degrees. Of note, the reaction with the D74A mutant exhibited the highest inhibitory effects, indicating that the salt bridge of D74 with R87 and the hydrogen bonds with R81’ and A84 were crucial for the pairing of the two proteins (Fig. 3A). Surprisingly, E55 forms multiple interactions with MazF-mt9, but its mutation barely affected the association of the TA pair (Fig. 5A and B).

**FIG 5.**
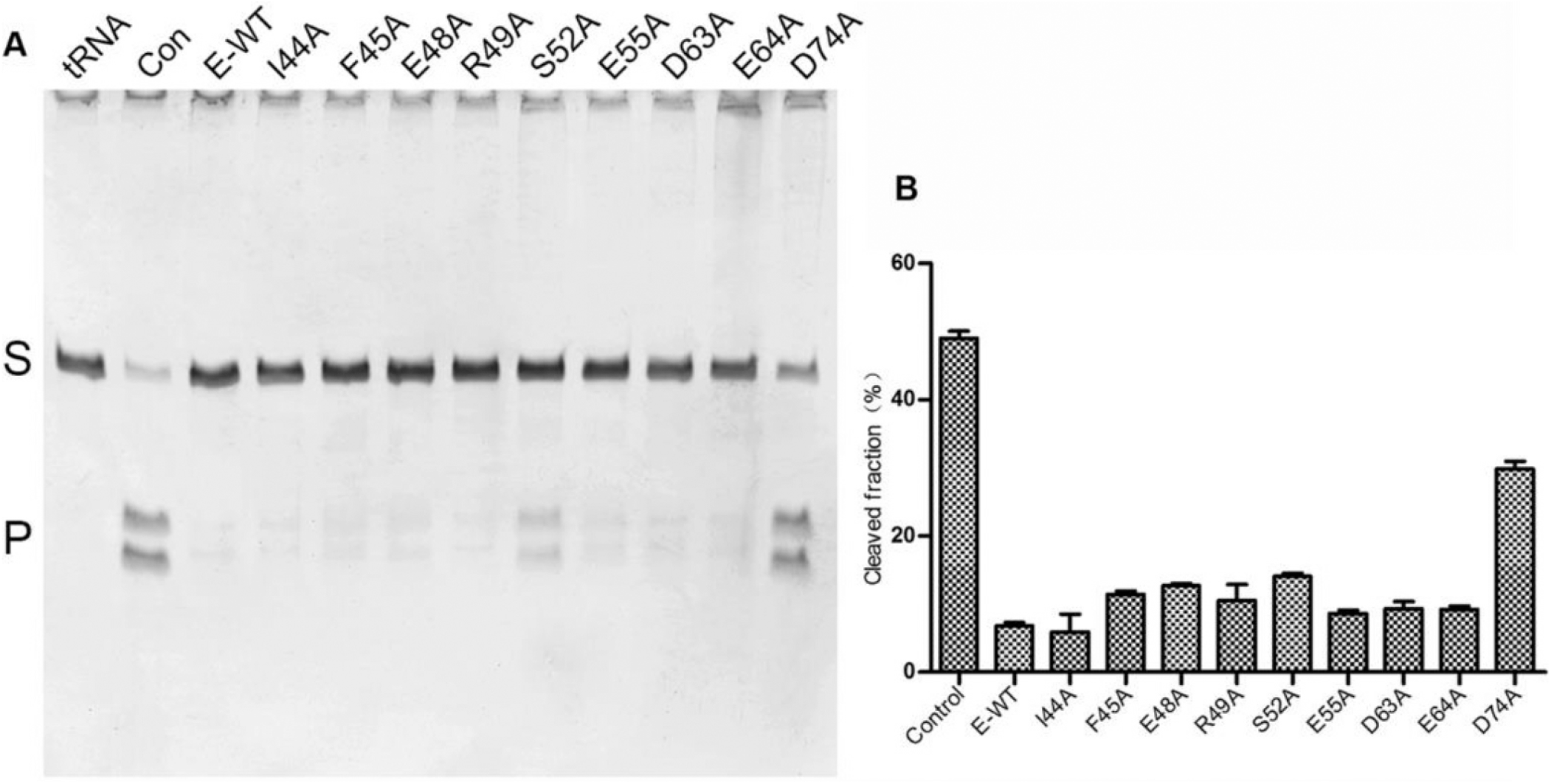
The activity cleavage tests on MazF-mt9 in complex with different mutants of MazE-mt9. Urea-PAGE gel (A) and summary by bar graphs (B) showing the cleavage results of MazF-mt9 in complex with different single mutants of MazE-mt9. Cleaved fraction was calculated by the ratio of the product intensity to the intensity of the starting substrate. The results were obtained from at least two independent reactions and error bars represent standard error of means from two independent experiments. Con: Control reaction by WT MazF-mt9 without the addition of the antitoxin. S: tRNA substrate. P: tRF products (2 bands).

We lastly measured the affinity of MazE-mt9 to MazF-mt9 by Isothermal titration calorimetry (ITC). The affinity of WT MazE-mt9 to MazF-mt9 is very tight, and *K*_D_ is in the nanomolar range (~17.5 nM) (Fig. 6A). The single D74A mutation approximately increased the *K*_D_ value by 2-folds (Fig. 6B), while the R46A-E48A-D74A triple mutation increased *K*_D_ by 8-folds (Fig. 6G). Furthermore, we removed the last helix containing D74, and this deletion mutant (SF62) completely lost its binding capability to MazF-mt9, underscoring the significance of D74 to the affinity between the MazE/F-mt9 complex. In contrast, other mutations did not cause significant changes to the *K*_D_ values (Fig. 6B-H).

**FIG 6.**
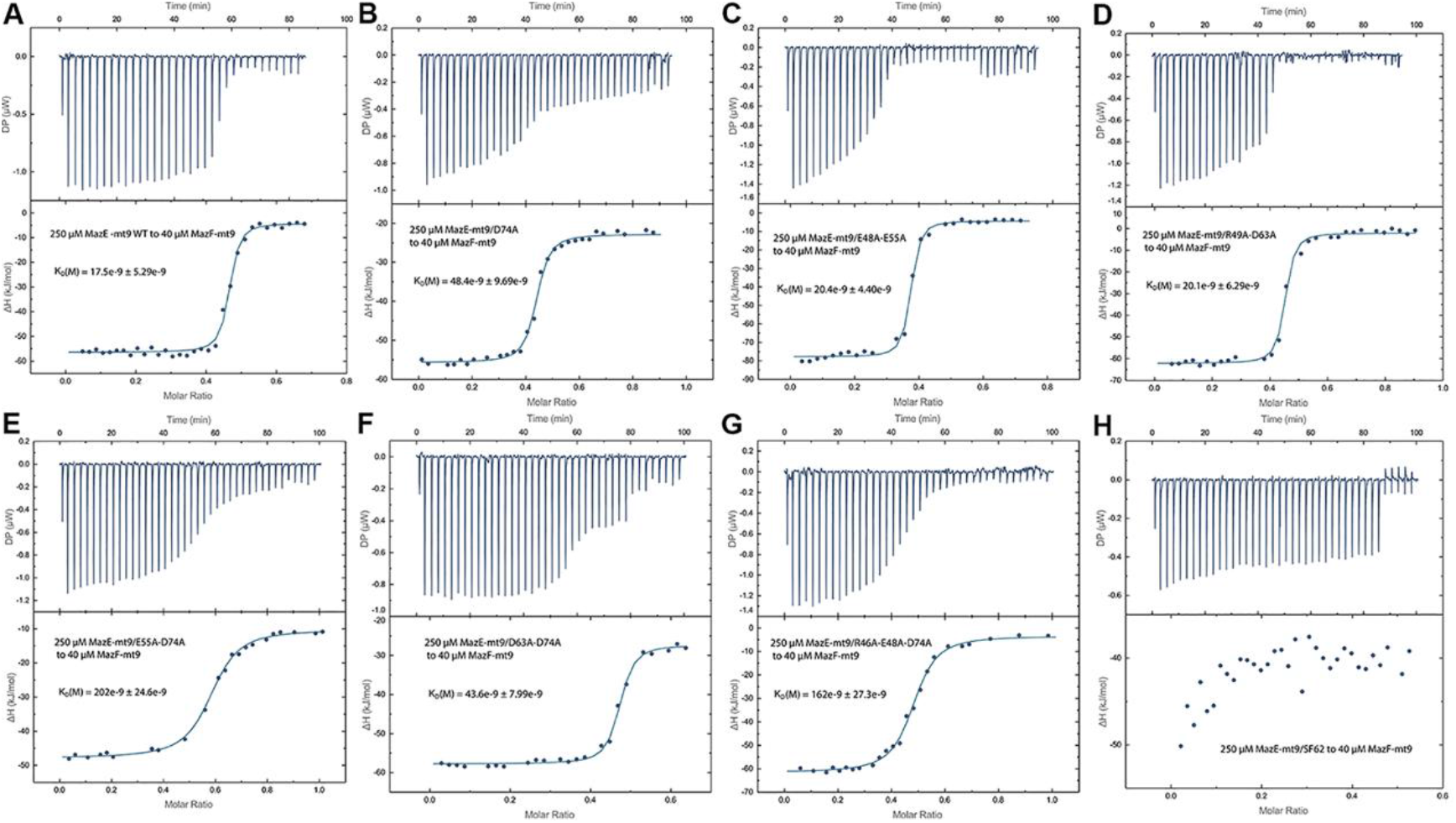
ITC evaluation on the contribution of the key residues in MazE-mt9 to the binding of the TA pair. (A)-(H) showed the binding profiles for the MazE-mt9/WT, MazE-mt9/D74A, E48A-E55A, R49A-D63A, E55A-D74A, D63A-D74A, R46A-E48A-D74A and SF62 mutants, respectively. The calculated *K*_D_ values are shown above the fitted curves.

### Characterization of the affinity between heterologous MazE/F complexes

*M. tb* contains ten MazF families. It is yet to be determined whether the MazF-mt9 family binds to other MazE members across the families. Although previously MazE-mt1 has been shown to bind MazF-mt3 (25), it is not known whether it was an individual case or a commonly occurring event. Structural analyses revealed that α3 of MazE-mt9 was key to the interactions with MazF-mt9. Additionally, secondary structure predication suggested that both MazE-mt1 and MazE-mt3 adopted a similar topology to that of MazE-mt9. Additionally, sequence alignment indicated that the conservation with MazE-mt3 mostly occurs at the N-termini of both proteins whereas the conservation with MazE-mt1 is mostly observed at the C-termini (Fig. S1). Therefore, MazE-mt1 would bind MazF-mt9 with higher affinity than MazE-mt3. As indicated by Fig. 7A and B, MazE-mt1 moderately reduced the cleavage activity of MazF-mt9 whereas MazE-mt3 hardly had any effects. Due to the weakened binding affinities of MazE-mt1 and -mt3, the unbound MazF-mt9 protein still showed “leaked” activities even the antitoxin is in excess. In addition, the D69A mutation of MazE-mt1 (equivalent of D74A on MazE-mt9) abolished its binding, as did the SF62 mutation or an irrelevant DNA-binding protein (Uracil-DNA glycosylase from *Nitratifractor salsuginis*, NsaUNG, Fig. 7B). These findings indicated that the activity of MazFs could be differentially antagonized by MazEs from different families.

**FIG 7.**
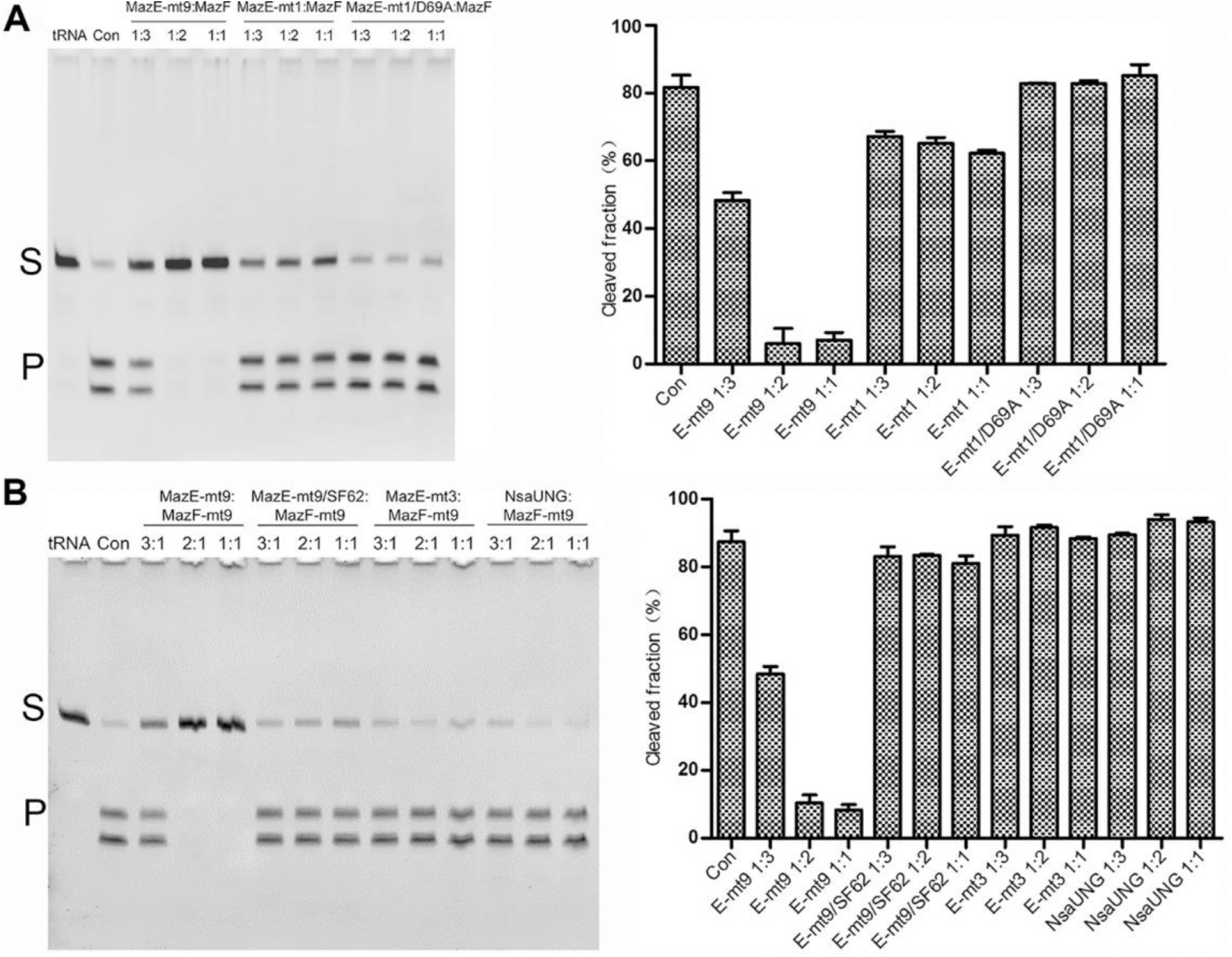
Interactions of MazF-mt9 with heterologous MazE members as assessed indirectly by cleavage activity assays. (A)-(B): Cleavage assays of MazF-mt9 in the presence of MazE-mt1 or the D69A mutant (A); or with MazE-mt3, MazF-mt9/SF62 or NsaUNG (B). S: tRNA substrate. P: tRF products (2 bands). The cleavage tests by each heterologous pairs were performed at three molar ratios: 1:3, 1:2, and 1:1 (toxin:antitoxin).

We next tested the *in*-*vivo* interacting abilities between the heterologous pairs through the co-expression of *maz*E-mt1/*maz*F-mt9 and *maz*E-mt3/*maz*F-mt9. The former resulted in the over-production of both the toxin and antitoxin molecules, and the protein pair could be co-purified through the N-terminal 8 × His-tag fused to MazE-mt1, suggesting stable interactions between them. These interactions, however, could be effectively abolished by the D69A mutation in MazE-mt1, where the induction of the pair harboring the mutation only generated the D69A mutant of the antitoxin (Fig. 8A and B). In contrast, co-expression of *maz*E-mt3 with *maz*F-mt9 did not produce significant amounts of either protein (data not shown). ITC results showed that the *K*_D_ value for the MazE-mt1/MazF-mt9 pair to be 59.3 nM, which explained the relatively strong inhibitory effect of MazE-mt1 against MazF-mt9 activity (Fig. 8C). By contrast, no measurable interactions between the non-cognate MazE-mt3/MazF-mt9 pair were detected by ITC (Fig. 8D).

**FIG 8.**
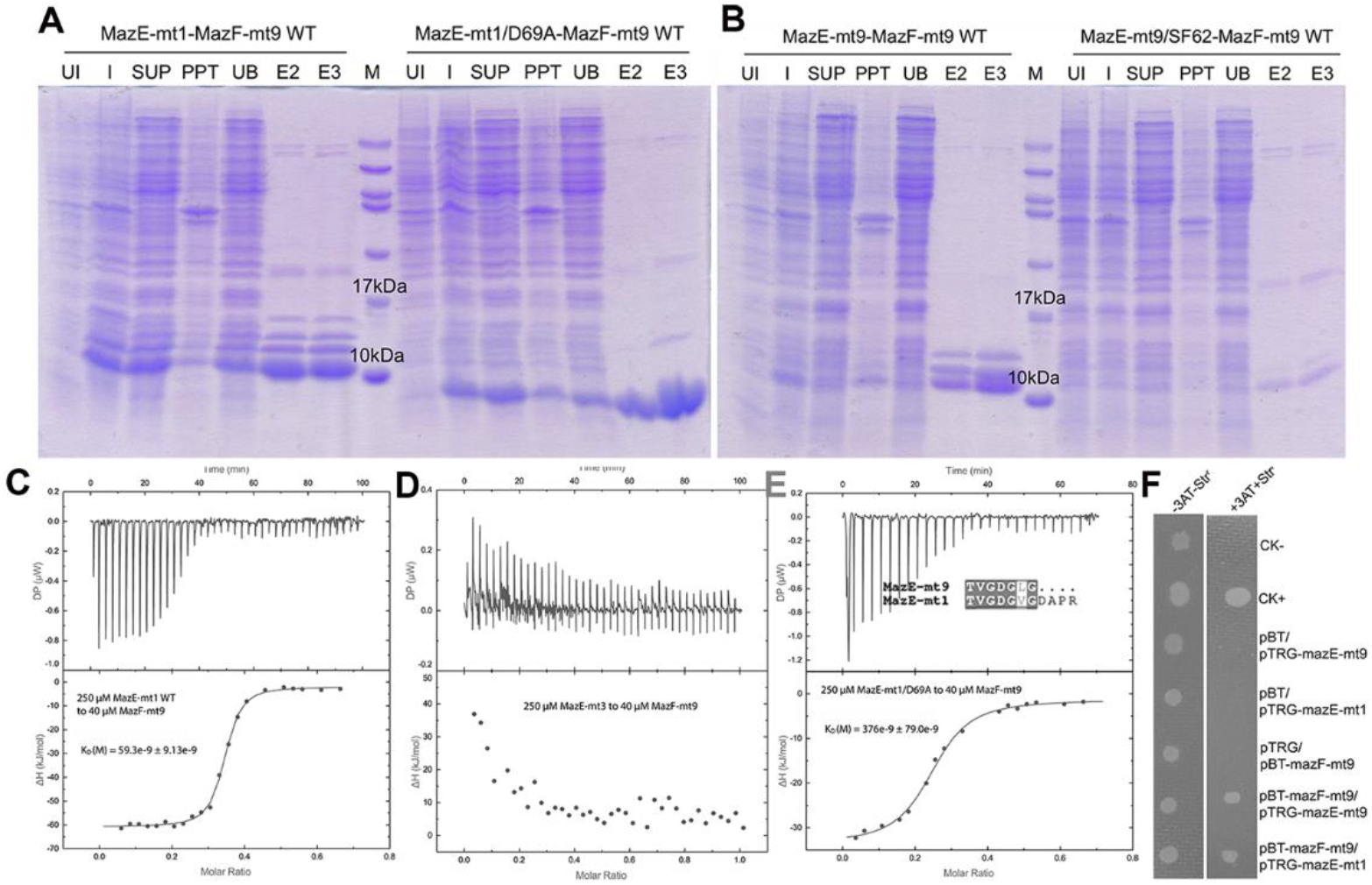
Further characterizations on the *in-vitro* and *in-vivo* interactions of MazF-mt9 with heterologous MazE members. (A) The co-expression and Ni-NTA purification of the heterologous MazE-mt1/MazF-mt9 complex, as visualized by SDS-PAGE electrophoresis. The profile for the MazE-mt1/MazF-mt9 complex was shown on the left in (A), whereas the MazE-mt1/MazF-mt9 complex with the D69A mutation of MazE-mt1 was shown on the right in (A). (B) The same as in (A) except that MazE-mt1 was replaced by MazE-mt9. (C-E) ITC measurements of the association affinity of MazF-mt9 with MazE-mt1 (C), or with MazE-mt3 (D), or with the MazE-mt1/D69A mutant. (F) The verification of the *in-vivo* MazF-mt9/MazE-mt1 interactions by the bacterial two-hybrid assay with (right lane) or without (left lane) 3-AT and streptomycin.

Furthermore, the conserved residues between MazE-mt1 and MazE-mt9 are concentrated at their C-termini. Of note, the TVGDGL(V) motifs are almost completely conserved (Fig. 8E). Importantly, the key aspartate residue for interactions could be found both in MazE-mt1 (D69) and MazE-mt3 (D80). Coincidentally, there are similar interactions in the BsMazEF pair, where E79 of MazE forms a salt bridge with R81 and a hydrogen bond with R80’ backbone (PDB 4ME7) (21). In addition, E80 in EcMazE makes identical interactions to EcMazF (R86 and K79), neutralizing the positive charges that otherwise would become buried upon EcMazE binding (PDB 5CQX) (26). ITC showed that the D69A substitution weakened the affinity of the non-cognate MazE-mt1 to MazF-mt9 by 5.3-folds, confirming the pivotal role of this aspartate (Fig. 8E). Consequently, the α-helices located at the C-termini of MazEs with the key aspartate residue may constitute the structural basis for the heterologous toxin-antitoxin interactions. Taken together, these findings indicated that the strongest interactions were best achieved by cognate MazEF pairs.

### Confirmation of non-orthogonal interactions in vivo

To further confirm our observation of the non-cognate interactions *in vivo*, we performed the bacterial two-hybrid assay, which is similar to the well-known yeast two-hybrid assay except for the choice of different host systems. The detection of protein-protein interactions depends on transcriptional activation of the *his*3 reporter gene. The expression of *his*3 allows *E. coli* growth in the presence of 3-amino-1,2,4-triazole (3-AT), a competitive inhibitor of the *his*3 enzyme, and positive co-transformants are verified by streptomycin resistance conferred by the *aad*A gene. The co-transformants expressing *mazF*-mt9/*mazE*-mt9 or *mazE*-mt1 were plated, and colony growth would indicate the existence of interactions between the protein pairs. As expected, both the cognate and non-cognate co-transformants grew well on the selective screening medium, meaning that MazF-mt9 indeed interacted with MazE-mt9 and MazE-mt1 respectively. On the other hand, while the pBT/*LGF*2 and pTRG/*Gal*11^p^ positive co-transformants (CK+) grew well on the plate, which was a proof of the principle, the pBT and pTRG empty vectors serving as the negative control (CK-) did not grow at all (Fig. 8F). However, the colony sizes resulted from MazF-mt9 non-orthogonal interactions were smaller than those of the pBT/*LGF*2 and pTRG/*Gal*11^p^ positive co-transformants, suggesting that both the toxic and the antidotal effects exist *in vivo* and the two opposing forces played an important part in cell growth.

## DISCUSSION

*M. tb* is a severe pathogen that has been plaguing humankind for a long history. The bacterium harbors over 90 reported TA systems, which are very conserved among the *M. tb* species. Most prokaryotes have only 1–2 MazEF TA systems. However, 10 MazEF TA systems have been discovered in *M. tb* (16, 17), which is quite amazing. According to previous studies, not all MazEF TA systems work independently. In fact, the relationships between different MazEF TA systems are rather complex in that they have independent functions, but they can also work together toward a common goal. In this work, we studied the crosstalks between MazF-mt9 and MazEs both *in vitro* and *in vivo* and proved that the cognate and non-cognate interactions indeed occur for this peculiar MazF member. Both types of interactions are rather strong, as judged from the dissociative constants (several tens milimolar range). These tight affinities explained the observation that most mutations barely disrupted the interactions between the TA pair. It also suggested that *in vivo*, the MazF-mt9 would be quickly sequestered by cognate or even non-cognate antitoxins. However, there could be discrepancies between the measured intensity of these interactions under the two circumstances (*in vitro* and *in vivo*) and it appeared that the mutations consequences *in vivo* were more moderate than *in vitro*. These differences were reasonable because the *in-vitro* experiments (tRNA-cleavage tests, ITC etc.) employed simple systems where the factors and amounts of proteins were easily manageable. In contrast, the *in-vivo* environment is more complex and as an indirect means to measure TA interactions, what we observed could be a compound effect for assays like colony growth. There could be other factors such as backup pathways or other agonists/antagonists involved.

Most MazFs are nucleases, and their distinct substrate specificities allow them to act on all three major types of RNAs (19, 27–32). mRNA cleavage inhibits synthesis of a certain protein, and its impact is usually mild. Hence this type of translation inhibition is considered as a minor “setback” for cellular processes. On the other hand, tRNA cleavage leads to aminoacylation failure of a certain amino acid and will certainly interfere with the synthesis of most proteins. Therefore, the tRNAse activity of MazF-mt9 will cause major damages to cells. Worst of all, the degradation on rRNA of the ribosome affects the translation process of all proteins, which is the most serious consequence of all three types of RNA degradation.

Therefore, under most stressful conditions, many MazF members will work synergistically. For example, the mRNA levels of *maz*F-mt1, *maz*F-mt3 and *maz*F-mt6 have been reported to increase in the event of a variety of stresses such as oxidation, nitrification, starvation and antibiotics treatment, suggesting that different MazFs could respond to the same stress signal (18). Additionally, the differential changes on expression levels suggested different participation extent of each member. In another scenario, the knockout of all three, but not any of the single or double genes of *maz*F-mt1, -mt3, -mt6 enhanced the susceptibility of *M. tb* to antibiotics. These results suggested that MazEF TA systems could indeed coordinate with each other to effect *M. tb*’s survival under highly stressful conditions (18).

Previous work indicated that crosstalks exist not only between different MazEF systems, but also between families outside the systems. For example, MazF-mt3 is capable of interacting with not only MazE-mt1 (MazEF interactions), but also with VapB24 and VapB25 from the VapBC system (non-MazEF interactions) (25). Moreover, MazF-mt7 was found to interact with DNA topoisomerase I, a non-TA system protein (33). Our study indicated that MazF-mt9 not only bound MazE-mt9 but also MazE-mt1, but the cognate interactions were much stronger than the heterologous interactions. These interactions have been observed both *in-vitro* and *in*-*vivo* in our study. Because both antitoxins could exist *in vivo* simultaneously, MazE-mt1 may play a helper role in inhibiting the activity of MazF-mt9 when there is not enough MazE-mt9 around. Due to the complexity of ten MazEF systems present in *M. tb*, we could not check all the cross-interactions, but it is very likely that these crosstalks exist between different systems. Zhu *et al.* think that these crosstalks challenged the “one toxin-one antitoxin” paradigm (25). With more and more crosstalk interactions being revealed, we think that TA systems will form an “interactome” instead of the simple, idealistic “one-on-one” interactions held by traditional views. The wide cross interactions will presumably help the cells to recover rapidly from the persister condition once the stressful pressure is lifted. In this case, both the cognate and heterologous interactions work together to reduce their toxicities. We speculate that *in*-*vivo*, the cognate interactions are the prevalent forces whereas the non-orthogonal interactions serve a secondary or accessory role to help to reduce the amount of free MazF-mt9 and thus fine-tune the activity of this toxin.

## MATERIALS AND METHODS

### Cloning, expression, and purification of the proteins

The genes encoding *M. tb maz*E-mt1 (Rv2801A), -mt3 (Rv1991A) and -mt9 (Rv2063) were amplified by PCR from the genomic DNA of the *M. tb* H37Rv strain. After double digestion by the *Sac*I and *Sal*I restriction enzymes, the PCR products were ligated into multiple cloning site 1 (MCS1) of the modified pETDuet1 (Novagen), in which a 8 × His-tag and a PreScission protease cleavage site were placed immediately upstream of MCS1. The mutants were generated by the *QuikChange* method (Stratagene) using the WT genes as the template. The MazEF co-expression vectors (including the mutants) were created by inserting the *maz*F gene into MCS2 of the above-mentioned pETDuet1/*maz*E plasmid. All the plasmids were transformed into *E. coli* strain BL21 (DE3) cells, and the cells were cultured in Luria-Bertani broth containing 50 μg/ml ampicillin at 37°C. The overexpression was induced by 0.3 mM isopropyl β-D-1-thiogalactopyranoside (IPTG) when the OD_600_ value reached 0.6–0.8. Three induction temperatures were 18, 25 and 37°C for MazF-mt9, MazE-mt9 and the MazEF-mt9 complex respectively. The purification of the proteins involved a two-step strategy comprising the Ni-NTA affinity and anion exchange chromatography. The target proteins without tag were obtained by collecting the unbound fractions from the Histrap affinity column (GE Healthcare) after digestion with the PreScission protease overnight (mass ratio = 1:100). The proteins of interests were concentrated and flash-frozen before storage at −80°C.

### Crystallization, data collection, and structure determination

Initial crystallization screening was set up by a Mosquito crystallization robot (TTP Labtech) using the sitting-drop vapor-diffusion method at room temperature. To obtain the cocrystals of the MazEF-mt9 complex, the MazF-mt9 protein was incubated with MazE-mt9 at a molar ratio of 1:1 in a buffer containing 20 mM Tris-HCl (pH 8.0), 150 mM NaCl, and 1 mM DTT, and incubated on the ice for 2 h. The final protein concentrations of MazE-mt9 and MazF-mt9 in the complex were 0.8 mg/ml and 1.3 mg/ml, respectively. The sample of the complex was centrifuged at 23,500 g for 10 min prior to crystallization. Crystal screenings have been carried out using commercial kits including the Index, Crystal Screen I & II, PEG/ion screens (Hampton research). Hits were observed overnight, and both needle-and rod-shaped crystals were obtained, generated in conditions containing PEG3350 or ammonium sulfate as the precipitant respectively. After optimization, thick, long rod-shaped crystals were obtained from 1.3–1.5 M ammonium sulfate and 0.1 M MES (pH 5.5). The fully grown crystals were soaked in a freshly-made cryoprotective solution containing all the components of the reservoir solution plus 20% (v/v) glycerol. The soaked crystals were mounted on nylon loops and flash-cooled in liquid nitrogen.

A 2.7-Å diffraction dataset was collected using beamline17U (BL17U) at the Shanghai Synchrotron Radiation Facility (SSRF, Shanghai, P. R. China) and was processed with the program HKL2000. Cell content analysis suggested that the asymmetric unit contained two MazE-mt9 and two MazF-mt9 molecules. The structure was solved by molecular replacement using the program *PHENIX* with the coordinates of *M. tb* MazF-mt9 structure (PDB 5WYG) (24) as the search model. The initial model was based on the solution and built manually according to the electron density map with *COOT* (34). Multiple cycles of refinement alternating with model rebuilding was carried out by *PHENIX.refine* (35). The final model was validated by *molprobity* (36). The structural figures were produced with *PyMOL* (www.pymol.org). All data collection and refinement statistics are presented in Table 1.

### ITC measurements

ITC experiments were conducted at 25°C using a PEAQ ITC titration calorimeter (Malvern instruments). To exactly match their buffer compositions, the MazE and MazF proteins (WT or mutants) were dialyzed against the same buffer containing 20 mM Tris-HCl (pH 8.0), 150 mM NaCl, and 1 mM DTT. The affinities were measured by titrating MazE or its mutants (250 μM) into MazF (40 μM). The first injection of 0.5 μL was followed by 19 injections of 2-μL drops. The MICROCAL ORIGIN software was used to determine the site-binding models that produced good fits. Individual peaks from titrations were integrated and displayed on a Wiseman plot. The first reading was removed from the analysis. The binding affinity (*K*_D_) and change in enthalpy (ΔH) associated with the binding events were calculated after fitting the dataset.

### In-vitro cleavage assays by the MazEF-mt9 complex

The preparation of the *M.tb* tRNA^Lys(UUU)^ transcripts was described in a previous protocol (37). 25 pmol of MazE-mt9 was mixed with 50 pmol MazF-mt9 and incubated on ice for 30 min to allow the formation of the complex. 0.6 μL of the complex was added to a reaction mixture of 20 mM HEPES (pH 7.5), 50 mM potassium chloride, 1 mM DTT and 8 pmol of *M. tb* tRNA^Lys(UUU)^substrate, and were incubated at 37°C. The reactions were stopped after 30 min by adding the 2 × formamide gel-loading buffer (95% w/v formamide and 50 mM EDTA). The samples were denatured at 95°C for 5 min prior to electrophoresis in a 15% Urea-PAGE gel containing 7 M urea, followed by ethidium-bromide staining.

### The overexpression effects of MazF-mt9 or the TA pair on E. coli growth

A single colony of the *E. coli* strain BL21 (DE3) cells carrying the pETDuet1/*maz*E-mt9, pCold TF/*maz*F-mt9 or the pETDuet1/*maz*EF-mt9 co-expression plasmid was added into 5 ml of Luria-Bertani broth containing 50 μg/ml ampicillin and was allowed to grow with rotation at 37°C overnight. The next morning the saturated culture was diluted to OD_600_ = 1.0 with LB broth containing 50 μg/ml ampicillin, and inoculated with 20 ml of LB. The OD_600_ value was measured each hour after it reached 0.4. Alternatively, LB culture with an OD_600_ value of 1.0 was further diluted for 10-, 100-, and 1000-folds, and 0.5 μL of each was transferred to LB-agar plates containing ampicillin, which had been pre-treated with or without 0.1 mM IPTG. The growth of *E. coli* colonies at 37°C on these two types of plates were monitored the next day.

### Bacterium two-hybrid assay

The BacterioMatch II two-hybrid system (Stratagene) was based on the *his*3-*aad*A reporter cassette (38). The amplified *mazF*-mt9 gene was cloned into the *EcolR*I-and *Xho*I-digested pBT plasmid, and the similarly treated *mazE*-mt1 and *mazE*-mt9 genes were each cloned into the *EcolR*I- and *Xho*I-digested pTRG plasmid. Pairs of the bait and prey plasmids were co-transformed into the *E*. *coli* XL1-Blue MRF’ strain, and the co-transformants were selected on M9-medium plates containing kanamycin (30 μg/ml), chloramphenicol (34 μg/ml), tetracycline (12.5 μg/ml), and 3-AT (4 mM) and streptomycin (8 μg/ml). The plates were incubated at 37°C for 3–4 days. Co-transformants expressing pBT/*LGF*2 and pTRG/*Gal*11^P^ (Stratagene) were used as positive controls for expected growth on the selective screening medium, while those containing the empty vectors pBT and pTRG were used as negative controls.

## ACCESSION NUMBER

The atomic coordinates and structure factors have been deposited in the Protein Data Bank, www.wwpdb.org (PDB 6A6X).

## SUPPORTING INFORMATION

Supplemental material for this article may be found at mBio.

**FIG S1,** PDF file, 0.3 MB.

**TABLE S1,** PDF file, 0.1 MB.

**TABLE S2,** PDF file, 0.1 MB.

## ACKNOWLEDGEMENTS

This work was supported by Natural Science Foundation of China 31870782, Natural Science Foundation of Guangdong province 2018A030313313, and Major program of collaborative innovation on Guangzhou health and therapy 201604020019.

We thank the staff members from BL17U1 beamlines at Shanghai Synchrotron Radiation Facility for assistance during data collection.

WX conceived and designed research; RC, JT, XC, CY, XD, BS, SM, XL, PM, and CD performed research; WX, YT and JR analyzed data and wrote the paper. All authors reviewed the results and approved the final version of the manuscript.

---

**FIG S1.**
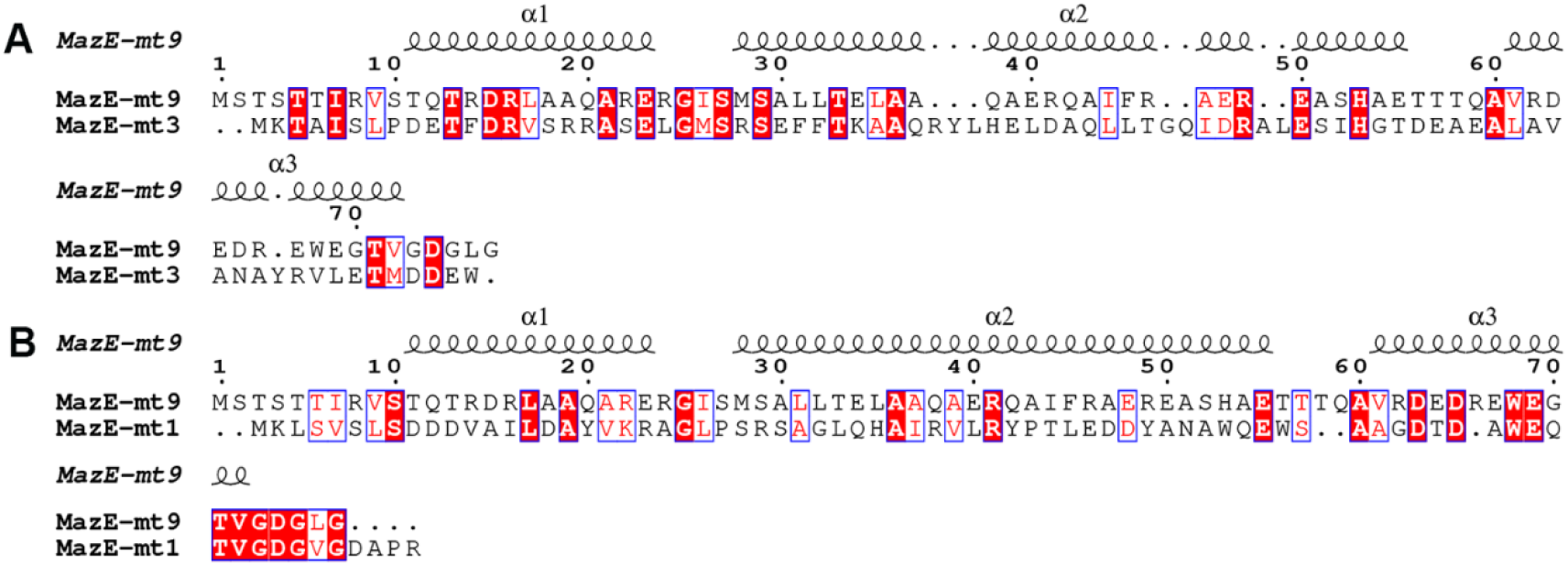
Sequence alignment of MazE-mt9. (A) Alignment with MazE-mt3, and with (B) MazE-mt1. The secondary structure of MazE-mt9 was shown above the alignments.

**Table S1.** The summary of the interactions within the MazEF-mt9 TA system.

| Chain C (MazE) | Chain A<br>(MazF) | Chain B<br>(MazF') |
| --- | --- | --- |
| I44 |  | P58/L59/L109 |
| F45 | V33/L36 | L109 |
| E48 |  | T57/R60 |
| R49 | E35/G-3 |  |
| S52 |  | R78 |
| E55 |  | R54/S55/R60 |
| D63 |  | S51 |
| E64 |  | R81 |
| D74 | R81 | R87 |

**Table S2.**
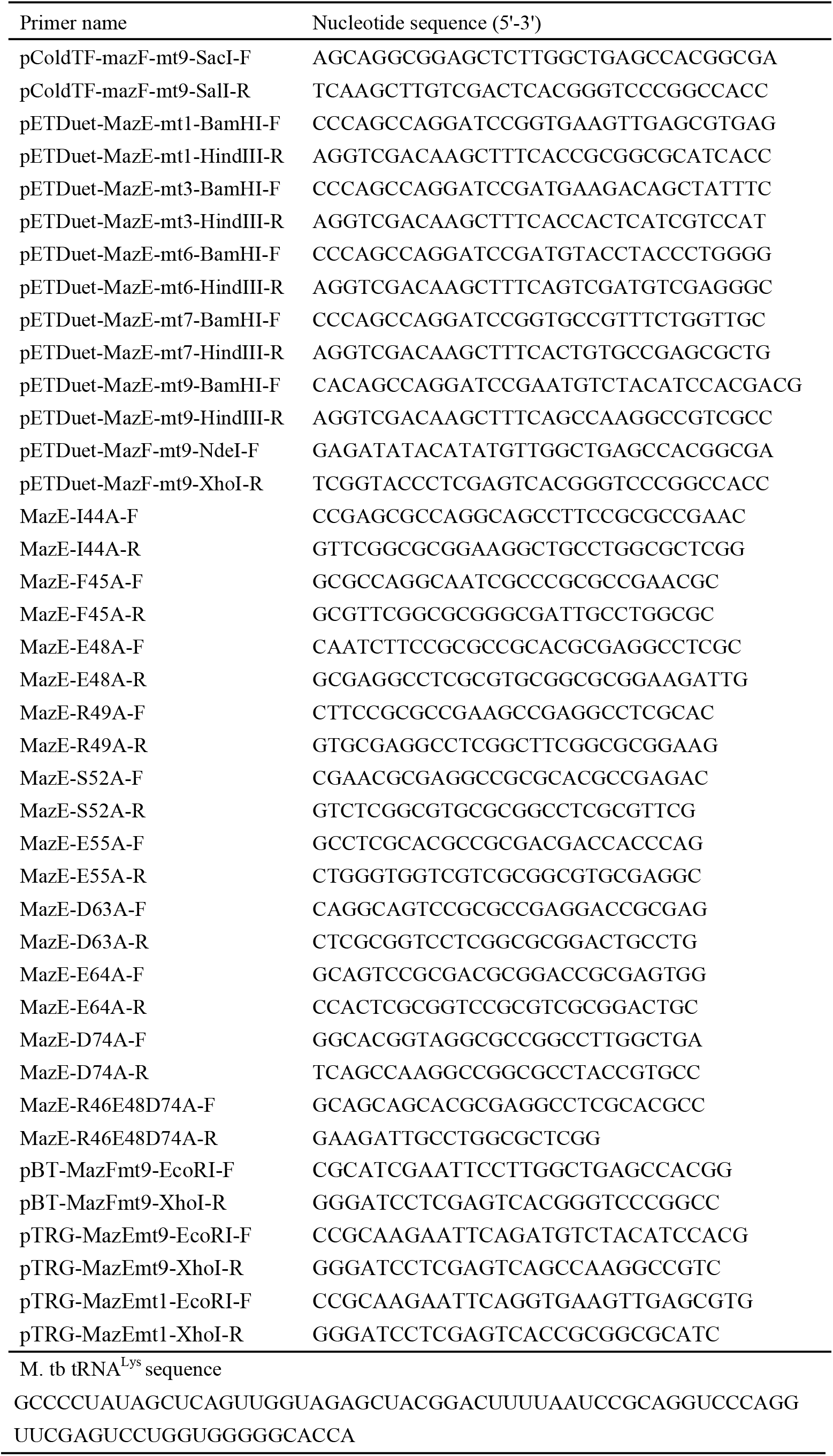
Sequences of the primers and tRNAs used in this study.

